# G6PD Inhibition Sensitizes Ovarian Cancer Cells to Oxidative Stress in the Metastatic Omental Microenvironment

**DOI:** 10.1101/2022.04.20.488962

**Authors:** Shree Bose, Qiang Huang, Yunhan Ma, Lihua Wang, Grecia O. Rivera, Yunxin Ouyang, Regina Whitaker, Rebecca A. Gibson, Christopher D. Kontos, Andrew Berchuck, Rebecca Previs, Xiling Shen

**Author notes:** Correspondence to: Xiling Shen.

## Abstract

Ovarian cancer (OC) is the most lethal gynecological malignancy, with aggressive metastatic disease responsible for the majority of ovarian cancer related deaths. In particular, OC tumors preferentially metastasize to and proliferate rapidly in the omentum. Here, we show metastatic OC cells experience increased oxidative stress in the omental microenvironment. Metabolic reprogramming, including upregulation of the pentose phosphate pathway (PPP), a key cellular redox homeostasis mechanism, allows OC cells to compensate for this challenge. Inhibition of G6PD, the rate-limiting enzyme of the PPP, reduces tumor burden in pre-clinical models of OC, suggesting this adaptive metabolic dependency is important for OC omental metastasis.

**Highlights:**

- The omental microenvironment poses a high oxidative stress metastatic niche for ovarian cancer cells.
- G6PD, a key enzyme involved in redox homeostasis and the rate-limiting enzyme of the pentose phosphate pathway (PPP) is upregulated in omental metastasis.
- Inhibition of G6PD increases oxidative stress and cytotoxicity in the omental microenvironment.
- Pharmacological G6PD inhibition reduces omental metastases *in vivo*.

## Introduction

Ovarian cancer (OC) remains the most lethal gynecological malignancy, with over 80% of patients diagnosed at Stage III or higher, when 5-year survival rates are ∼30% (Matulonis et al., 2016). An overwhelming majority of OC deaths are attributed to metastatic disease, with exfoliation from the primary tumor into the peritoneal fluid leading to peritoneal dissemination of cancerous cells throughout the abdominal cavity (Lengyel, 2010). The most clinically relevant and common site of metastatic growth is the omentum, an apron-like layer of adipose tissue which lays over the peritoneal cavity. Omental tumors often grow quite aggressively, leading to significant consequences for patient morbidity and mortality. The precise molecular mechanisms which drive this aggressive phenotype of OC cells in the omental microenvironment are not yet understood but are thus of particular interest.

Alterations in metabolism have recently been proposed as a potential avenue of cellular reprogramming to promote ovarian tumor metastasis; however, the precise nature of these changes is not yet well understood. Evidence suggests that OC cells harness available fatty acid resources in the omental microenvironmental niche (Fidler, 2003; Lengyel, 2010; Nieman, 2011). Observed upregulation of proteins critical to cellular fatty acid (FA) metabolism, like fatty acid binding protein 4 (FABP4), a carrier protein to import free FAs, has provided critical support for this model (Gharpure, 2018; Ladanyi, 2018; Mukherjee et al., 2020). In addition, adipocytes in the metastatic niche have also been seen to undergo cellular remodeling to support tumor growth and invasion—secreting adipokines to stimulate the adhesion, migration, and invasion of tumor cells, releasing local paracrine factors within the tumor microenvironment, and adapting lipid metabolism to provide free fatty acids for fuel (Lengyel et al., 2018; Nieman et al., 2013).

The shift towards OC fatty acid metabolism is accompanied by alterations in the omental stroma to create the metastatic niche. To support the increased FA uptake of metastatic OC cells, activated omental adipocytes release reactive oxygen species (ROS) like H_2_O_2_ and NO, increasing local oxidative stress (Herroon et al., 2018). Canonically, the cellular antioxidant response is largely reliant on the critical reducing equivalent, nicotinamide adenine dinucleotide phosphate (NADPH). This cofactor is recycled between NADPH and its oxidized form NADP+, to serve as an electron donor for antioxidant defense and reductive biosynthesis. A key enzyme responsible for NADPH generation is glucose-6-phosphate dehydrogenase (G6PD), the rate-limiting enzyme of the oxidative pentose phosphate pathway (PPP). This parallel pathway to the glycolytic machinery has been uniquely implicated in as a requirement for NADPH/NADP+ redox homeostasis, mammalian dihydrofolate reductase (DHFR) activity, and folate metabolism in cancer cells (Chen et al., 2019). While G6PD is a potential mediator of chemotherapeutic response (Feng et al., 2020; Yamawaki et al., 2021); the potential role of this redox enzyme in OC omental metastases has not been elucidated.

Here, we investigated alterations in G6PD as an OC adaptation in the omental microenvironment. Both human and mouse omental metastases and *in vitro* models of omental metastases of OC exhibited higher oxidative stress and upregulated G6PD. Inhibition of G6PD led to increased cell death in the omental microenvironment *in vitro* and reduced metastases *in vivo*. Thus, G6PD may play a role in mitigating oxidative stress and promoting growth in OC omental metastases.

## Results

### Omental metastases exhibit upregulated G6PD expression and activity in the context of higher oxidative stress

OC cancer cells exhibit a specific tropism and rapid growth rate in the metastatic niche of the omentum. We first studied the transcriptomic profiles of 30 primary ovarian tumors and 30 matched omental metastases from patients with Stage III high grade serous ovarian adenocarcinomas using available NCBI Gene Expression Omnibus datasets (GEO: GSE118828 and GSE30587). Hierarchical clustering and t-SNE plotting (Figures S1A and 1A) suggested distinct transcriptomic profiles for primary tumors vs. omental metastases. Paired differential (Figure 1B) and pathway analysis (Figure 1C) of a panel of metabolic genes obtained from the KEGG Database (http://www.kegg.jp) was conducted, and the PPP emerged as an altered pathway in metastatic samples.

**Figure 1:**
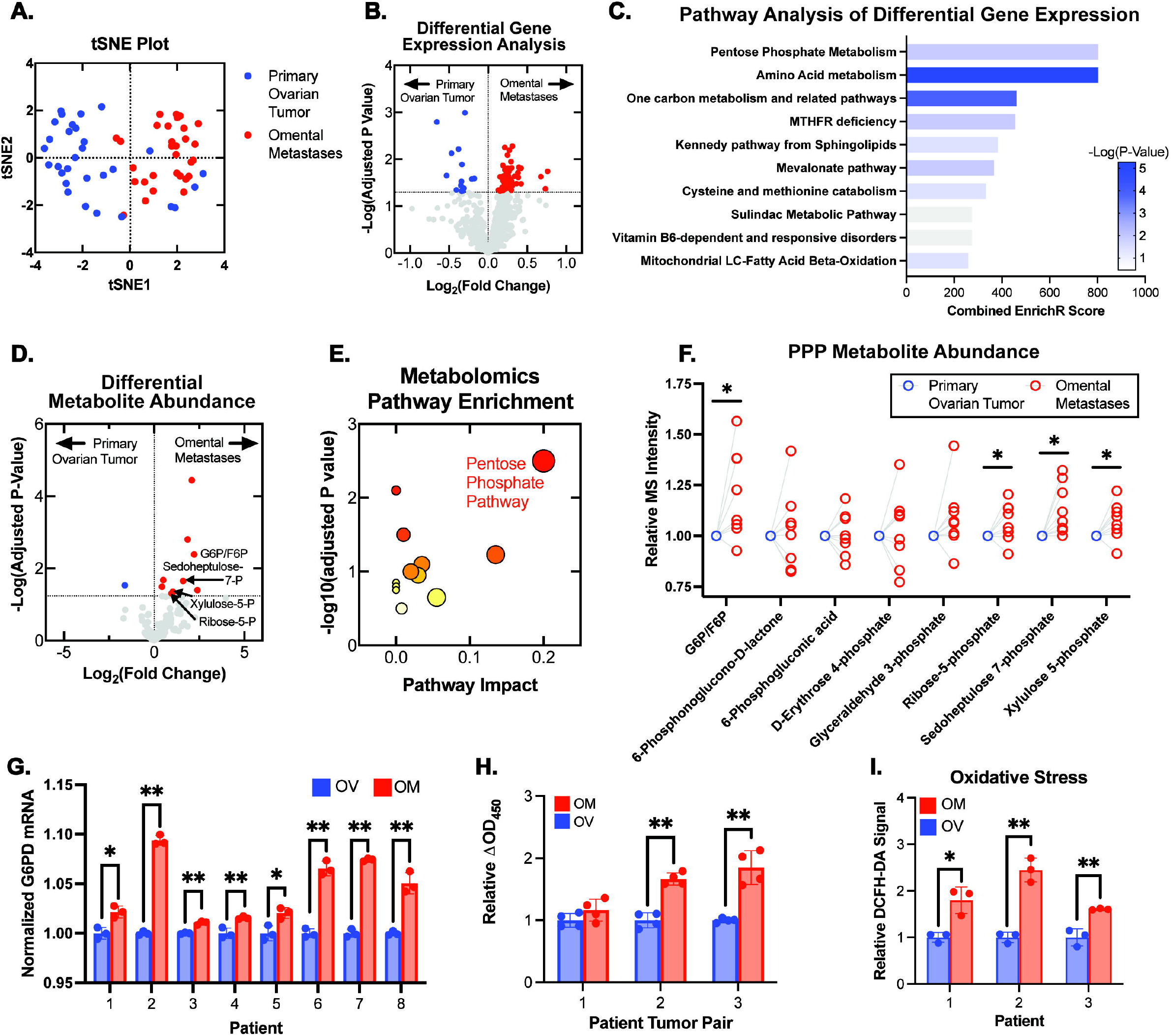
The pentose phosphate pathway is upregulated in omental metastases compared to primary ovarian cancers. Gene expression data (n=30) revealed metabolic differences of primary ovarian tumors and omental tumors from Stage III OC patients. **(A)** Distinct clustering in tSNE plot of metabolic gene data from matched primary ovarian tumors (blue) and omental metastases (red). **(B)** Volcano plot of gene expression changes for paired ovarian and omental tumors highlighted genes significantly upregulated in omental tumors (red) and ovarian tumors (blue). **(C)** Pathway analysis of the significantly altered genes. **(D)** Volcano plot of metabolite abundances determined by LC-MS metabolomic analysis of matched ovarian and omental pairs (n=8). **(E)** Pathway enrichment analysis of metabolomics data; symbol size indicates pathway impact and color indicates p-value. **(F)** Changes in PPP metabolite abundances are plotted as outlined circles. Blue circles indicate the normalized primary tumor measurements for all tumor pairs (n=8). **(G)** G6PD expression in ovarian and omental metastases (n=8) was quantified using qPCR. **(H)** G6PD activity of tumor pairs (n=3) was measured via enzymatic assay. **(I)** Tumor lysates (n=3 pairs) was incubated with 25 µM DCFH-DA and fluorescence was measured at 30 minutes. Statistical significances are noted as * p<0.05, ** p<0.01 by two-tailed Student’s *t*-Tests.

To further characterize the phenotype, we performed metabolomic profiling of matched primary ovarian tumors and omental metastases from eight patients with Stage III/IV high grade serous ovarian cancer (HGSOC). A targeted analysis reported relative quantities of 124 polar metabolites, and differential analysis of metabolite abundances was performed (Figure 1D). Pathway and enrichment analysis of the statistically significant hits identified alterations in PPP metabolism (Figure 1E and S1B). PPP metabolites emerged as highly differential between primary ovarian tumors and the corresponding omental metastasis (Figure 1F).

In matched patient primary OC-omental metastasis pairs, expression of G6PD, the rate-limiting enzyme of the PPP responsible for generating the key reducing equivalent NADPH, was increased in omental metastases, measured using qPCR (Figure 1F) and immunoblotting (Figure S1C and S1D). G6PD activity measured using an enzymatic assay was similarly increased in omental metastases (Figure 1H). We then used dichlorodihydrofluorescein diacetate (DCFH-DA), a cell-permeable fluorometric probe, to evaluate levels of intracellular reactive oxygen species (ROS) in the matched pairs. DCFH-DA assay measurements indicated higher intracellular ROS in extracts from omental metastases compared to primary tumors in these samples (Figure 1I).

Using an established model of murine ovarian cancer metastases to replicate the features of primary ovarian tumors and omental metastases, we performed intraperitoneal injections of each of three common high grade serous ovarian cancer (HGSOC) cell lines—HEYA8, SKOV3, and IGROV1—in immunodeficient mice (Shaw et al., 2004). These cell lines were transduced using lentivirus to stably express mCherry and luciferase. IVIS imaging of the xenografted nude mice at humane endpoints prior to sacrifice revealed omental metastases present in all cell line models (Figure 2A). To confirm, we used intravital imaging to follow metastatic seeding in the omentum after intraperitoneal injection of HEYA8-mCherry labeled cells. At 6 hours post intraperitoneal injection, disseminated micrometastases were visible in the mouse omentum; by 42 hours, growth into larger nodules was seen (Figure 2B).

**Figure 2:**
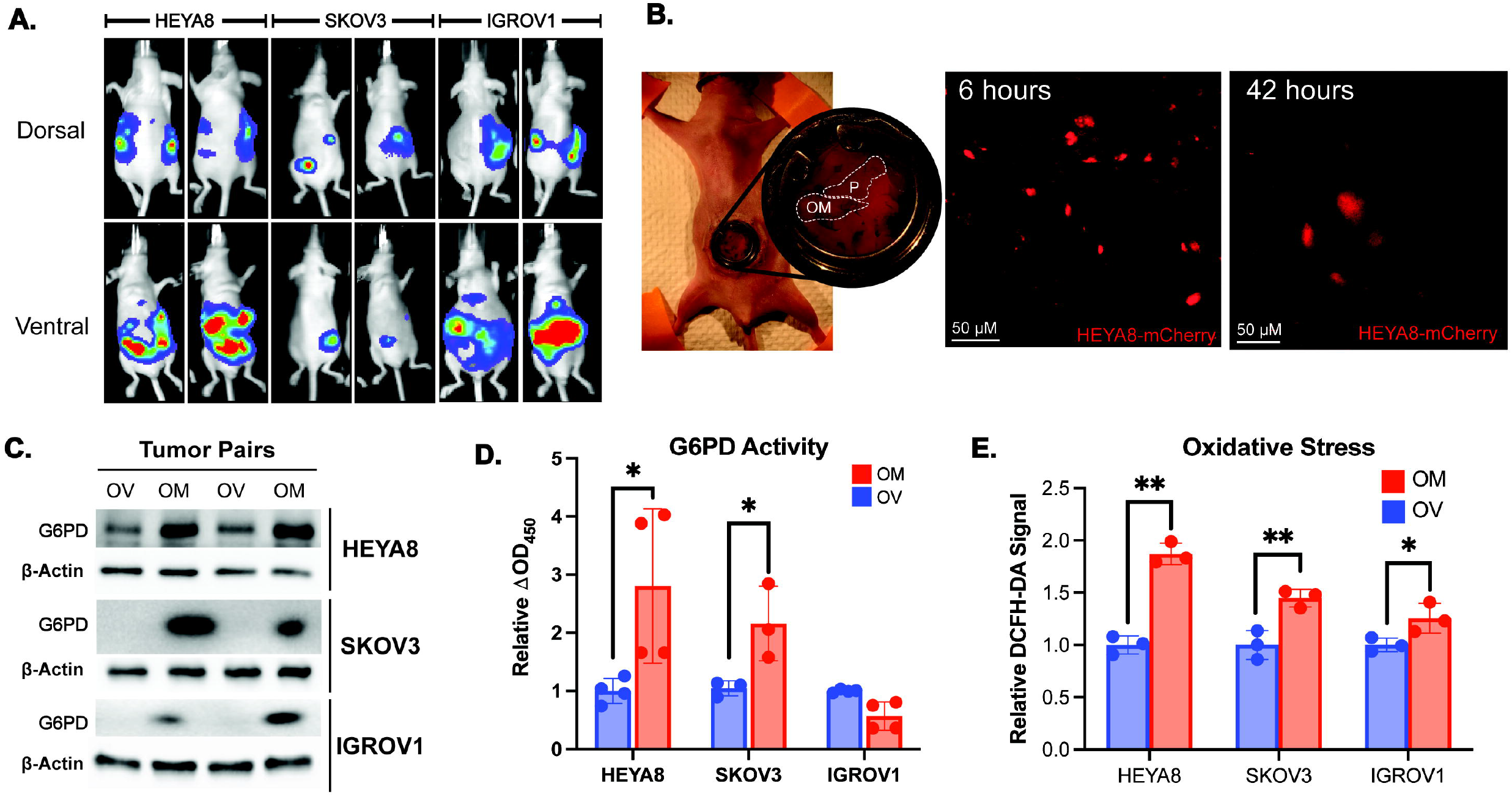
Murine models of ovarian cancer exhibit omental metastasis. **(A)** HEYA8, SKOV3, and IGROV1 tumors were imaged by IVIS. **(B)** Intravital imaging showed metastatic seeding and growth in the omentum at 6 and 42 hrs. **(C)** Immunoblotting for G6PD expression in murine omental tumor samples. **(D)** Enzymatic measurement of G6PD activity in primary tumors and omental metastases. **(E)** Tumor lysates were incubated with 25 µM DCFH-DA and fluorescence was quantified. Statistical significances are noted as * p<0.05, ** p<0.01 by two-tailed Student’s *t*-Tests.

WBs revealed increased G6PD expression in omental metastases (Figures 2C), consistent with our previous observation from human samples (Figures 1G). Enzymatic assays showed increased G6PD activity in omental metastases of HEYA8 and SKOV3, but not in IGROV1, which did not exhibit preferential metastasis to the omentum (Figure 2D). DCFH-DA assays showed higher oxidative stress in omental metastases, congruent with the human samples as well (Figure 2E).

Tumor organoids were also derived from the ovarian primary tumor and from the omental metastases of the three different murine models (Figure S2A). CellROX oxidative stress assays did not show significant differences in oxidative stress between the ovarian and omental tumor organoids cultured in the same organoid media composition (Figure S2B), suggesting that oxidative stress is likely driven by omental microenvironmental factors.

### Omental Conditioned Media Increased Oxidative Stress and G6PD Upregulation

The use of omental conditioned media (OCM) has been extensively validated and characterized in the literature as a model of omental metastases (Clark et al., 2013). We cultured OC cells in media conditioned with omental adipose tissue. Using DCFH-DA as a readout of intracellular ROS, oxidative stress in the OC cells grown in DMEM + 10% FBS was compared to those grown in OCM (Figure 3A). Both HEYA8 and SKOV3 showed higher oxidative stress in OCM-grown conditions, while IGROV1 remained relatively unchanged.

**Figure 3:**
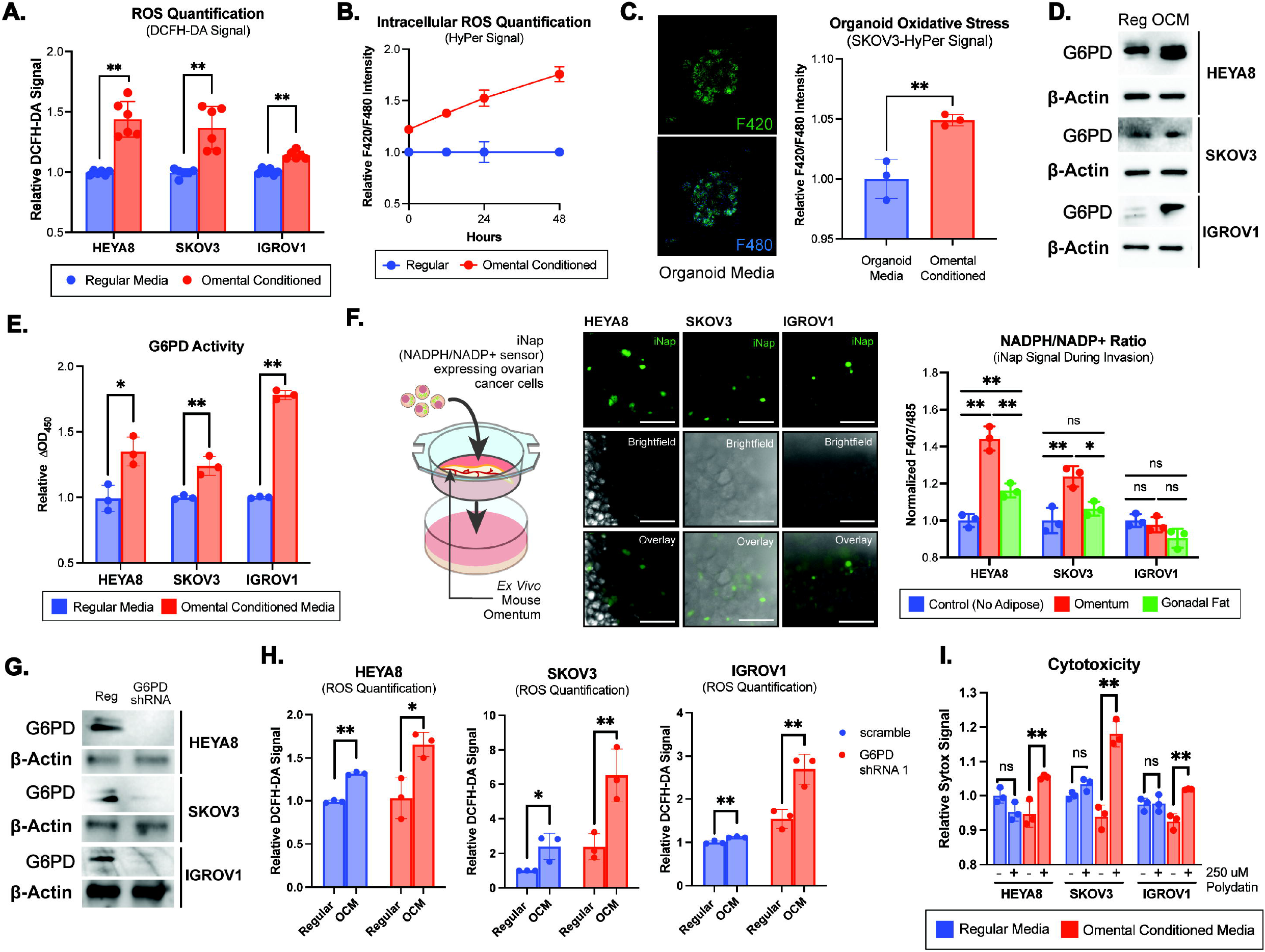
G6PD inhibition sensitizes ovarian cancer cells to the increased oxidative stress of the omental microenvironment. **(A)** OC cell lines grown in OCM were incubated with DCFH-DA and fluorescence was quantified. **(B)** SKOV3 OC cells were transfected with constructs for the intracellular H_2_O_2_ sensor, HyPer. Ratiometric fluorescence was measured at 0, 12, 24, and 48 hrs. **(C)** HyPer signal quantification of OC organoids cultured in omental conditioned organoid media. **(D)** G6PD expression via immunoblotting and **(E)** G6PD activity measured via enzymatic assay in OCM-cultured cells. **(F)** NADPH/NADP+ ratios of OC cells invading an *ex vivo* omental culture measured by iNAP, a NADPH/NADP+ ratiometric biosensor. **(G)** Immunoblotting for G6PD in OC cell lines after lentiviral shRNA knockdown. **(H)** Quantification of oxidative stress measured via DCFH-DA signal in wildtype and G6PD shRNA expressing OC cells cultured in OCM. **(I)** Quantification of cytotoxicity measured via Sytox staining of OCM and polydatin-treated OC cells. Statistical significances are noted as * p<0.05, ** p<0.01 by two-tailed Student’s *t*-Tests.

Given the rapid dynamics of the redox landscape and multiple layers of cellular regulation, we next sought to characterize oxidative stress using genetically, encoded fluorescent biosensors, which could more precisely quantify and track endogenous reactive oxygen species (ROS) dynamics. These include rapid interchanges of superoxide anions [O_2_·^−^] and hydrogen peroxide (H_2_O_2_) and reducing equivalents like NADPH. We first utilized HyPer, a genetically encoded, fluorescent biosensor of intracellular H_2_O_2_ to study OC cells cultured in OCM. After first establishing that HyPer signal was indeed responsive to oxidative stress induced by exogenous H_2_O_2_ (Figure S2C), we used a lentiviral vector to produce a SKOV3 cell line that stably expresses the HyPer biosensor. When grown in OCM, HyPer signal increased over time, indicating higher oxidative stress in the cells (Figure 3B). Furthermore, to account for potential difference between 2D and 3D culture, the biosensor-expressing cells were grown in 3D Matrigel to form OC organoids for dynamic metabolic analysis. Using ratiometric excitation at 420 and 480 nm with emission readings at 500 nm, fluorimetry was used to determine dynamic changes in oxidative stress. SKOV3-HyPer organoids exhibited higher oxidative stress when exposed to omental conditioned organoid culture media (Figure 3C).

HEYA8, SKOV3, and IGROV1 OC cells grown in OCM showed higher expression and activity levels of G6PD (Figure 3D and 3E). Furthermore, 10 OC cell lines reflected differing degrees of OCM-induced oxidative stress increases and G6PD activity increases (Figure S3A and S3B), suggesting G6PD may be employed as a part of the cellular response to oxidative stress.

A key aspect of G6PD’s role in redox homeostasis involves the generation of NADPH, an essential electron donor in all organisms and a critical reducing power for anabolic reactions and redox balance. In order to determine if the process of metastasis into the omentum affected the cellular balance between NADPH and its oxidized counterpart, NADP+, a modified *ex vivo* 3D coculture model was developed (Krishnan et al., 2015). OC cells expressing iNAP, a ratiometric NADPH/NADP+ biosensor described in the literature (Tao et al., 2017), were cocultured with an omental explant and the NADPH/NADP+ ratio was measured during the dynamic process of omental invasion. The NADPH/NADP+ ratio was markedly higher in HEYA8 and SKOV3 cells invading the omentum compared to simple migration (Figure 3F). Gonadal fat was also isolated and used as a control explant for invasion. OC cells invading omenta exhibited higher NADPH/NADP+ ratios than those invading gonadal fat, suggesting a greater need for NADPH production in omental-invading cancer cells.

### G6PD Inhibition Sensitized Ovarian Cancer Cells to Oxidative Stress

We next investigated if OCM-grown cells experienced higher oxidative stress when compensatory G6PD mechanisms were inhibited. We first established successful lentiviral shRNA knockdown of G6PD using WB (Figure 3G) and then evaluated intracellular ROS using DCFH-DA. In HEYA8, SKOV3, and IGROV1 cells, shRNA knockdown of G6PD increased oxidative stress, with a combination of growth in OCM and shRNA knockdown is responsible for higher oxidative stress than baseline (Figure 3H).

Polydatin, a pharmacological inhibitor of G6PD (Mele et al., 2018), was then used to treat OC cells grown in regular media and OCM. Cytotoxicity was significantly increased in OCM-cultured cells treated with polydatin as compared to normal culture conditions (Figure 3I). Measures of cellular viability including Crystal Violet and the formazan-based WST-8 assay reflected similar trends, with less viable cells evident in OCM-polydatin treated samples (Figure S3C and S3D). In cells treated with polydatin, increased NADPH/NADP+ ratio was also noted (Figure S3E). Treatment with a known ROS scavenger, N-acetylcysteine (NAC), reversed these cytotoxic effects *in vitro*, validating that increased oxidative stress caused by OCM growth was indeed responsible for the cell death seen in polydatin-treated cells (Figure S3F).

### G6PD Inhibition Reduces Omental Metastases *In Vivo*

Having established that G6PD inhibition could sensitize cells to OCM-induced oxidative stress, we next sought to explore its effect on omental metastases *in vivo*. HEYA8-mCherry-luciferase cells stably expressing G6PD knockdown shRNA were injected into NSG mice. Evaluation of tumor burden using IVIS imaging revealed reduced tumor growth in the G6PD-knockdown cohort (Figure S4A and S4B). At sacrifice, isolation of peritoneal fat deposits and subsequent fluorescence imaging of the mCherry-expressing tumors revealed greater omental tumor burden in the mice injected with G6PD-expressing HEYA8 cells than G6PD-knockdown HEYA8 cells (Figures S4C). Metastatic tumor growth in the gonadal and mesenteric fat were not significantly different between the two cohorts (Figure S4D and S4E).

To study the effects of pharmacological G6PD inhibition via polydatin on OC tumor growth and metastasis *in vivo*, we then injected athymic, nude mice with HEYA8 ovarian cancer cells labeled with mCherry and luciferase tags. Initial tumor engraftment was not significantly different between the treatment and vehicle cohorts (Figure S5A and S5B). Daily doses of 100 mg/kg polydatin were injected intraperitoneally, and a control group of mice were injected with vehicle as shown in Figure 4A. Tumor growth was followed longitudinally throughout the study using IVIS imaging, during which total bioluminescence (BLI) was quantified (Figure 4B). No adverse effects were noted with polydatin, and weights remained consistent between the vehicle and treatment cohorts (Figure S5E). To evaluate if polydatin treatment affected growth of metastatic tumors in the omentum, mice were imaged and sacrificed 14 days after treatment initiation. IVIS imaging revealed reduced overall tumor burden in the polydatin-treated cohort (Figures 4C and 4D) at time of sacrifice. Upon gross dissection, primary tumors and associated peritoneal metastases were significantly larger in the vehicle-treated mice than those treated with polydatin (Figure 4E). Isolation of the omentum and subsequent fluorescence imaging for the mCherry-expressing tumors revealed large tumors in the vehicle-treated omenta with only small puncta visible in the polydatin-treated (Figure 4F). Quantification of this difference revealed significantly reduced tumor growth in the omenta of polydatin-treated mice (Figure 4G). Staining of omental tumors was also performed and revealed similar tumor morphology between vehicle and polydatin treated tumors (Figure S5F), and a reduction in metastases to the mesenteric fat but not gonadal fat (Figure S5C and S5D). Treatment with 30 mg/kg polydatin did not appear to significantly decrease tumor burden at time of sacrifice nor metastases to the omentum, mesenteric, or gonadal fats (Figure S6).

**Figure 4:**
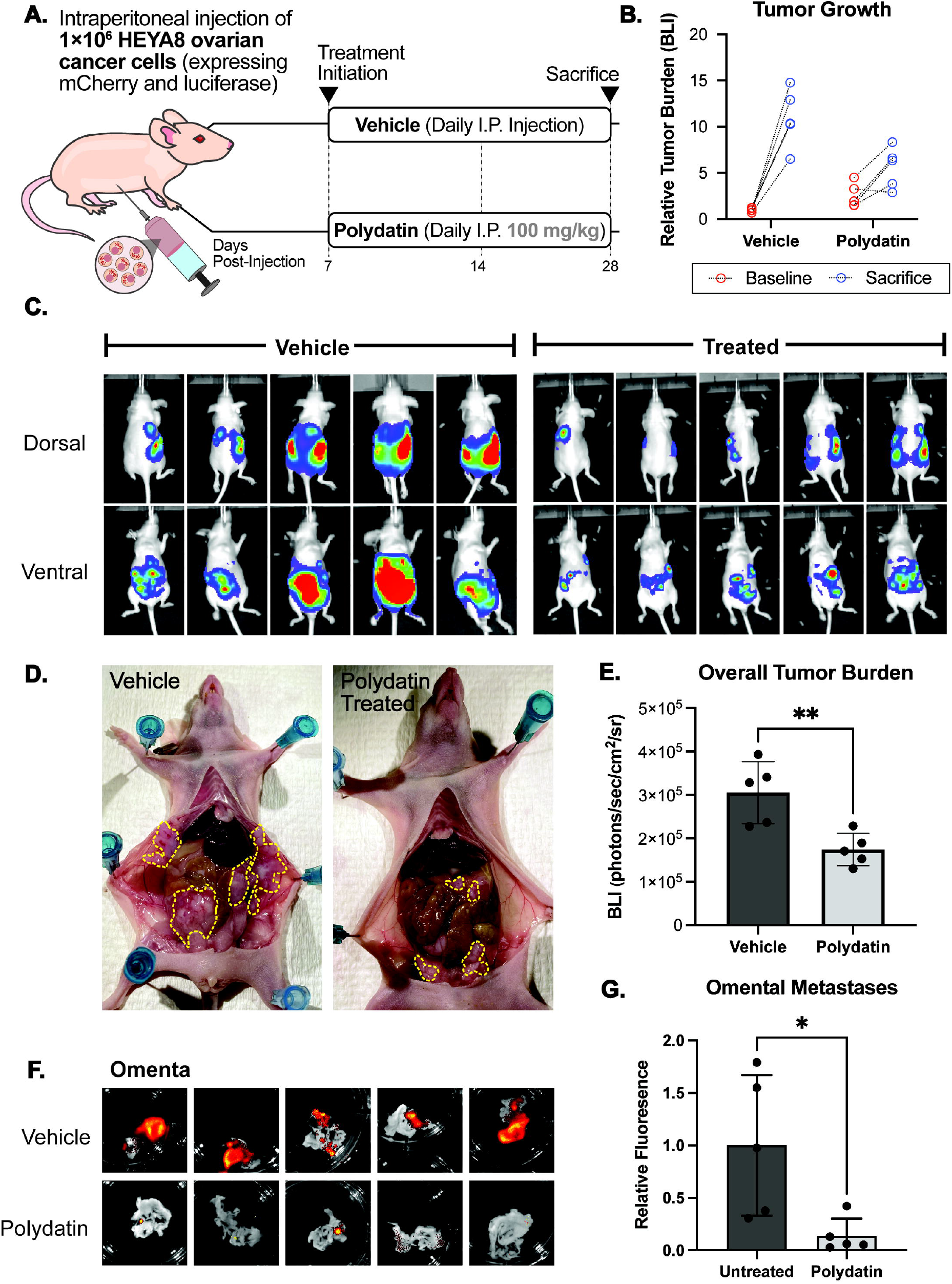
Polydatin treatment reduces omental metastases *in vivo*. **(A)** Schema for treatment. **(B)** Bioluminescence (BLI) was measured at baseline and prior to sacrifice using IVIS. Reduction in peritoneal metastases by polydatin treatment as measured by **(C)** IVIS imaging and **(D)** gross inspection of peritoneal tumor burden at necropsy. **(E)** Overall tumor burden as measured as BLI on IVIS in polydatin and vehicle cohorts (**p-value<0.01). **(F)** Fluorescence imaging and **(G)** quantification of omental metastases (*p-value<0.05).

## Discussion

The Warburg Effect is a well-known adaptation of cancer that cancer cells preferentially perform aerobic glycolysis in lieu of the more energy-efficient oxidative phosphorylation. Further studies have revealed other pathways of metabolic reprogramming in cancer, including amino acid and lipid metabolism (Beloribi-Djefaflia et al., 2016; Faubert et al., 2020). As the scope and complexity of our understanding of tumor metabolism expands, important transformations between primary and metastatic tumor metabolism emerge as critical differentiators and barriers to metabolic drug development that would effectively target both. Thus, a deeper understanding of the microenvironmental responses that both drive and enable survival of metastatic cells is particularly valuable. With the advent of new biochemical tools, deeper explorations of the role of oxidative stress in promoting or challenging neoplastic growth may reveal novel avenues to study and potentially target cancer (Hayes et al., 2020).

Here, we report an important role for G6PD in OC metastases to the omentum. While G6PD expression and activity in primary tumors has recently emerged as an area of interest in other cancer types, the unique metabolic reprogramming of the omental microenvironment—with increased local oxidative stress from immune infiltrates and adipose tissue FA synthesis—requires OC cells to compensate effectively. We found OC cells in this environment exhibit a dependency on G6PD, perhaps as a mechanism to provide this compensation, with shRNA knockdowns inducing significantly higher stress in the OC environment, and pharmacological inhibition using polydatin increasing cytotoxicity. *In vivo*, treatment with polydatin significantly reduced metastatic tumor burden in peritoneal fat deposits, including the omentum.

As G6PD is canonically considered an NADPH-producing enzyme, oxidative stress compensation may play a more important role in the metastatic process than previously appreciated in OC. The precise causality of this transformation—whether microenvironmental oxidative stress creates a demand for NADPH, simultaneously triggering FA metabolism or vice versa—is not well-understood, and likely dependent on multiple cellular processes. Indeed, OCM-induced oxidative stress and G6PD activity are heterogenous in human, murine, and *in vitro* OC models, implicating multiple cellular mechanisms. In this study, we have focused on understanding the role of the PPP, a redox homeostasis mechanism potentially employed by metastatic, omental OC cells as the demand to neutralize ROS outpaces other NADPH-producing mechanisms.

The importance of the omental microenvironment in driving an aggressive transformation of OC metastases is of clinical relevance, given the burden of OC metastatic disease on patient outcomes and recurrent disease. The reasons underlying the observed increases in oxidative stress responses seen in metastatic cells are likely manifold and will require further studies to fully characterize. G6PD, a key cellular redox homeostasis enzyme, may play an important role in this response, as inhibition of this protein affected OC growth in the omental microenvironment both *in vitro* and *in vivo*. Thus, our studies suggest this unique metabolic state caused by the omental microenvironment poses a context-dependent vulnerability which warrants further investigation, particularly as a potential therapeutic target.

## Limitations of the Study

The precise mechanism of polydatin has not been well-established, and whether inhibitory effects on G6PD mediates observed cardioprotective activity, antiinflammatory, immunomodulatory, and antioxidative effects remains unclear (Du et al., 2013). In addition, the high polydatin dose used in the animal experiments (100 mg/kg daily) may pose barriers for clinical application in humans. Future investigations using other inhibitors, including the estrogen precursor, Dehydroepiandrosterone (DHEA), the monocarboxylic acid amide, 6-aminonicotinamide (6-AN), or the more specific nonsteroidal inhibitor, G6PDi-1, would be of interest (Ghergurovich et al., 2020).

## Methods

### Human Studies

Human tissues were obtained from Duke University School of Medicine Ovarian Cancer Research Biobank under protocol ID Pro00013710 (Banking Normal and Malignant Gynecologic Tissues Removed at Surgery). Eight patients with Stage III/IV HGSOC were identified and corresponding primary ovarian tumors and omental metastases were obtained. Tumor samples were isolated at the time of tumor resection and debulking surgery, and snap-frozen in liquid nitrogen at time of collection. For assays, tissue was ground using mortar and pestle while maintained in liquid nitrogen, and ∼20 mg of tissue was used for DCFH-DA assays, G6PD enzyme activity assays, and WBs.

For metabolomics analysis, metabolite extraction from ground tissue normalized by weight was performed in cold extraction buffer (40% methanol: 40% acetonitrile: 20% water) solution. Extraction was performed by vortexing and centrifugation at 16,000 x g for 10 min, after which the supernatant was isolated and 15% NH_4_HCO_3_ was added to neutralize acids and precipitate proteins. Following centrifugation, the supernatant was applied to a speed vac and a concentrated metabolite pellet was obtained. Following reconstitution, the sample was used for LC-MS performed at the Rutgers Metabolomics Core Service.

### Animal Studies

Athymic, nude (Foxn1^nu^) female mice, 6-8 weeks old were obtained from the Jackson Laboratory and maintained in compliance with Institutional Animal Care and Use Committee (IACUC) guidelines. Mice were injected with 1×10^6^ ovarian cells intraperitoneally (n = 5 mice/group) for drug treatment studies or in the flank for organoid development. Where indicated, polydatin treatment (100 mg/kg) or 90% PBS + 10% DMSO (as vehicle) was administered daily for 14 days. Bioimaging was performed at different time points/once per week starting one week post tumor engraftment using an *in vivo* IVIS (IVIS Kinetic). Peritoneal metastases were evaluated based on mCherry signals by an OV100 microscope (Olympus) after sacrifice and dissection. All studies were performed in compliance with institutional guidelines under an IACUC-approved protocol. Tumor specimens were isolated and either snap-frozen for further analysis or processed for immunostaining or organoid development.

### Cell Culture Procedures

HEYA8, SKOV3, and IGROV1 ovarian cancer cells were cultured in DMEM medium supplemented with 10% fetal bovine serum (FBS) (Corning Life Sciences) and 1% penicillin/streptomycin/L-glutamine (Corning Life Sciences). Cells were periodically screened for Mycoplasma contamination.

Constructs for HyPer-DAAO (Thomas Michel, Addgene) and iNAP (Yiping Yi, gift), two genetically encoded metabolic biosensors were transfected using Lipofectamine 2000 and were maintained in DMEM + 10% FBS until fluorescent expression was seen under a fluorescence microscope (Olympus OV100).

Polydatin (Sigma-Aldrich) was dissolved in DMSO and aliquoted to stock solutions of 250 mM kept at -20, made fresh prior to cell culture experiments. Omental conditioned media was generated by isolating fresh omentum from sacrificed C57BL/6J mice. After 3 washes in PBS to remove residual clots from dissection, omenta were cultured in DMEM without FBS or penicillin/streptomycin for 48 hours. The adipose tissue was then removed, and the media was passed through a 45 µM filter prior to use in cell culture applications.

### Lentiviral mCherry/HyPer-DAAO/Luciferase expression and shRNA Knockdown

To generate lentivirus, HEK293T cells were transfected with psPAX2 and pMDG.2 packaging plasmids and the lentiviral expression construct. Viral supernatant was collected after 48 and 72 hours and concentrated using LentiX concentrator (Takara Bio). This virus was used to transduce cells with Transdux (System Bio). To generate mCherry-luciferase expressing cell lines, cells were stably infected with lentiviral constructs and FACS was used to select based on mCherry-expression. To generate HyPer-DAAO expressing cell lines, HyPer-DAAO was cloned into a lentiviral vector backbone prior to lentiviral generation using HEK293T cells. Bacterial glycerol stocks of G6PD shRNAs in a pLKO-puromycin backbone were obtained from Mission (Millipore Sigma) and were used to generate lentivirus. Following infection of OC cells, selection was performed using 5 ug/mL puromycin.

### Organoid Cultures

In brief, organoids were established according to previously described methods. Following isolation, tumor tissue was stored in AdDF+++ (Advanced DMEM/F12 containing 1x Glutamax, 10□mM HEPES and antibiotics), prior to being washed 3x in ice-cold PBS. Tissue was then minced finely in a small amount of HBSS and centrifuged to form a cell pellet. Digestion was carried out using AdDF+++ supplemented with 5□µM RHO/ROCK pathway inhibitor (Abmole Bioscience, Y-27632) containing 0.5–1.0□mg/mL collagenase (Sigma, C9407) at 37□°C for 1 hour, after which cells were spun down. The remaining cell pellet was resuspended in Matrigel (Corning Sciences, Inc.) and plated in 25 µL domes in 24 well plate. Organoid culture media composition is shown in Table S1.

### *Ex Vivo* Omental Invasion Assay

Using omenta isolated from sacrificed C57BL/6J mice, we performed an *ex vivo* assay as previously described (Krishnan *et al*., 2015). In short, after washing the omentum in HBSS, a small amount of Matrigel was applied to the bottom of a transwell plate and the omentum was gently placed upon it. Similarly, a thin strip of gonadal fat was used as a control and placed on the bottom of a transwell insert in a similar fashion. After 10 min at 37 degrees, DMEM + 10% FBS + 1% penicillin/streptomycin was applied to the bottom chamber, and serum-free DMEM was applied to the top chamber. OC cells expressing a fluorescence biosensor were applied to the top chamber and the fluorescence was quantified using a plate reader (Varioskan and Incucyte imaging during the invasion process.

### Computational Analysis

Two NCBI Gene Expression Omnibus (GEO) databases (GEO: GSE118828 and GSE30587) were identified that contain transcriptomic profiles of 30 primary ovarian tumors and 30 matched omental metastases from patients with Stage III high grade serous ovarian adenocarcinomas. High throughput RNA sequencing data was normalized to the primary tumor means and log2 transformed. Microarray expression data was analyzed using the Bioconductor package in R Studio, and normalized data was pooled for metanalysis. Hierarchical clustering was performed. The 20 patients with the greatest variation between transcriptomic profiles were identified by calculations of Euclidean distance between vectors of gene expression in omental metastasis and primary tumors. This cohort was used to perform pathway analysis in EnrichR (R package).

### Statistical Analysis

Statistics were performed using GraphPad Prism7 (GraphPad Software) or R and are specified in each figure legend. All data were tested for normality. Paired and unpaired two-tailed Student’s *t*-tests were used as appropriate for each experiment. A p-value of < 0.05 was considered significant, the level of significance is indicated as follows: ∗ = p < 0.05, ∗∗ = p < 0.01; n.s. indicates no significance. All data are presented as mean ± SD.

### Reagents and biochemical methods

#### DCFH-DA Assay

Within cells, the acetate groups of DCFH-DA are first cleaved by cellular esterizes to produce the intermediate DCFH, which then undergoes two electron oxidation reactions by intracellular ROS to generate the fluorogenic molecule DCF (Aranda et al., 2013). Cells were incubated in serum-free DMEM containing 10 µM DCFH-DA solution for 30 minutes prior to fluorescence readings at an excitation wavelength of 485 nm and an emission wavelength of 530 nm.

#### G6PD Activity Assay

The G6PD enzyme activity (Sigma Aldrich) was performed as described previously (Jiang et al., 2011; Tian et al., 1998).

#### Cytotoxicity/Viability Assays

Cells were plated at appropriate confluence in a 96-well plate (Corning Life Sciences), and allowed to seed for 24 hours, after which media was aspirated and replaced with appropriate treatment media. For viability assays, cells were allowed to grow for 48 hours, after which media was again aspirated and cells were incubated with Crystal Violet stain for 30 minutes. Stain was removed and wells were washed with PBS until only adherent cells appeared stained under light microscopy. Plate was allowed to air dry and appropriate solubilization agent was added, after which absorbance at 570 nm was read (Varioskan). For cytotoxicity assays, cells were allowed to grow for 72 hours, after which media was aspirated and cells were incubated with 2.5 µM Sytox Blue Reagant dissolved in PBS for 30 minutes at RT. Fluorescence was measured at excitation/emission of 405 nm/488 nm (Varioskan).

#### Immunostaining

Immediately following dissection, tumors were isolated and washed in PBS, prior to immersion in 30% sucrose overnight. After fixation in 4% neutral-buffered formalin, tumors were then embedded in Tissue-Tek® O.C.T. Compound (Sakura Finetek) and sectioned with a cryostat. H&E staining as previously described (Fischer et al., 2008) was then performed. Tumor sections were also immersed in PBST with 0.1% anti-G6PD antibody

#### Quantitative PCR

qPCR experiments were performed as previously described. Briefly, human tissue samples were ground using mortar and pestle in liquid nitrogen, and RNA was isolated using RNeasy kit (Zymo) and converted to cDNA using RevertAid First Strand cDNA Synthesis Kit (Thermo Fisher). We performed qPCR with a Taqman Gene Expression Assay using a QuantStudio 12K Flex machine (Applied Biosystems). Primers were obtained from ThermoFisher for Actin (Catalog #4331182) and G6PD (Catalog #4331182).

#### Western Blotting

Immunoblotting experiments were performed as previously described. Briefly, whole-cell lysates were generated by lysing cells in 50 mM Tris pH 7.5, 150 mM NaCl, 0.5% NP40, 1 mM EDTA, 10% glycerol, protease inhibitor mix (Roche) and phosphatase inhibitor cocktail (Sigma). Total protein amount was measured using a BCA assay, after which protein loading was normalized across samples. A Varioskan plate reader was used (Molecular Devices, Sunnyvale, CA, USA). Rabbit anti-G6PD antibody was obtained from Abcam, and mouse anti-rabbit HRP secondary antibody was obtained from Cell Signal.

## Acknowledgements

This work was supported by National Cancer Institute grants NIH-U01CA217514 and U01CA214300, as well as National Institutes of Health F30 fellowship 1F30CA257365-01. We would like to thank Dr. Yiping Yi (East China University) for providing the NADPH /NADP+ sensor, iNAP for use in this study, and Dr. Thomas Michel (Harvard Medical School) for providing the intracellular hydrogen peroxide HyPer-DAAO sensor. We would like to thank Regina Whitaker and the Duke OB/GYN clinical team for their assistance obtaining clinical samples. Services, results and/or products in support of the research project were generated by the Rutgers Cancer Institute of New Jersey Metabolomics Shared Resource, supported, in part, with funding from NCI-CCSG P30CA072720-5923.

## Supplementary Figures

**Figure S1: Gene expression and metabolomics analysis highlight PPP changes in matched human ovarian and omental tumors. (A)** Hierarchical clustering of heatmap of metabolic gene expression data from ovarian and omental tumors (n=30) show clustering of primary tumors and metastatic tumors independently. **(B)** Pathway analysis of metabolite abundance differences from LC-MS metabolomics analysis of matched primary ovarian tumor vs. omental metastases (n=8) highlight the pentose phosphate pathway. **(C)** Immunoblotting for G6PD in ovarian and omental tumors (n=3). **(D)** Quantification was performed by normalizing protein expression to the loading β-actin control and the primary tumor expression. Omental metastases exhibited increased G6PD expression (p-val<0.05, Student’s *t*-Test).

**Figure S2: *In vitro* models of OC reflect changes in oxidative stress. (A)** Organoids were derived from murine ovarian tumors and omental metastases. **(B)** Oxidative stress of organoids measured using CellROX reveal minimal differences. **(C)** Fluorimetry of OC cells expressing HyPer was responsive to hydrogen peroxide exposure.

**Figure S3: OC Growth in OCM Affects Cellular Behaviors.** Growth in OCM induced increases in **(A)** oxidative stress as measured via DCFH-DA incubation and **(B)** G6PD activity as measured via enzymatic assay in 10 OC cell lines. **(C)** Crystal Violet and **(D)** WST-8 cell viability assays revealed less viable cells evident in OCM-polydatin treated samples. **(E)** iNAP fluorescence revealed increased NADPH/NADP+ in cells treated with polydatin. **(F)** N-acetylcysteine (NAC) treatment reversed these cytotoxic effects *in vitro*. Where indicated, statistical significance is noted using * p<0.05, ** p<0.01 by two-tailed Student’s *t*-Test.

**Figure S4: G6PD-shRNA knockdown reduces omental metastases *in vivo*. (A)** IVIS imaging and **(B)** quantification of tumor burden in NSG mice injected with wild-type and G6PD-shRNA expressing HEYA8-mCherry cells. Fluorescence imaging and quantification of mCherry-expressing tumor burden in **(C)** omenta, **(D)** gonadal, and **(E)** mesenteric fat deposits.

**Figure S5: High-dose polydatin treatment (100 mg/kg) reduces mesenteric metastases *in vivo*. (A)** IVIS imaging and **(B)** quantification of baseline tumor engraftment taken 3 days post-injection of HEYA8-mCherry injected mice. Fluorescence imaging and quantification of mCherry-expressing tumor burden in **(C)** gonadal and **(D)** mesenteric fat deposits. **(E)** Weights of mice did not vary significantly during course of treatment (each symbol is average of 5 mice per cohort, no statistically significant differences seen). **(F)** H&E, G6PD staining, and mCherry microscopy of omental tumors revealed similar tumor morphology between vehicle and polydatin treated tumors.

**Figure S6: Low-dose polydatin (30 mg/kg) treatment does not dose not achieve therapeutic threshold *in vivo*. (A)** Schema for treatment. **(B)** Bioluminescence (BLI) was measured at baseline and prior to sacrifice using IVIS. **(C)** IVIS imaging and BLI quantification prior to sacrifice shows tumor burden in 30 mg/kg polydatin treated and vehicle treated mice. Fluorescence imaging and quantification of mCherry-expressing tumor burden in **(D)** omenta, **(E)** gonadal, and **(F)** mesenteric fat deposits. Labels of n.s. indicate “not significant” (p-val>0.05)

