## Supplementary Figures for "G6PD Inhibition Sensitizes Ovarian Cancer Cells to Oxidative Stress in the Metastatic Omental Microenvironment"

Supplementary Figure 1

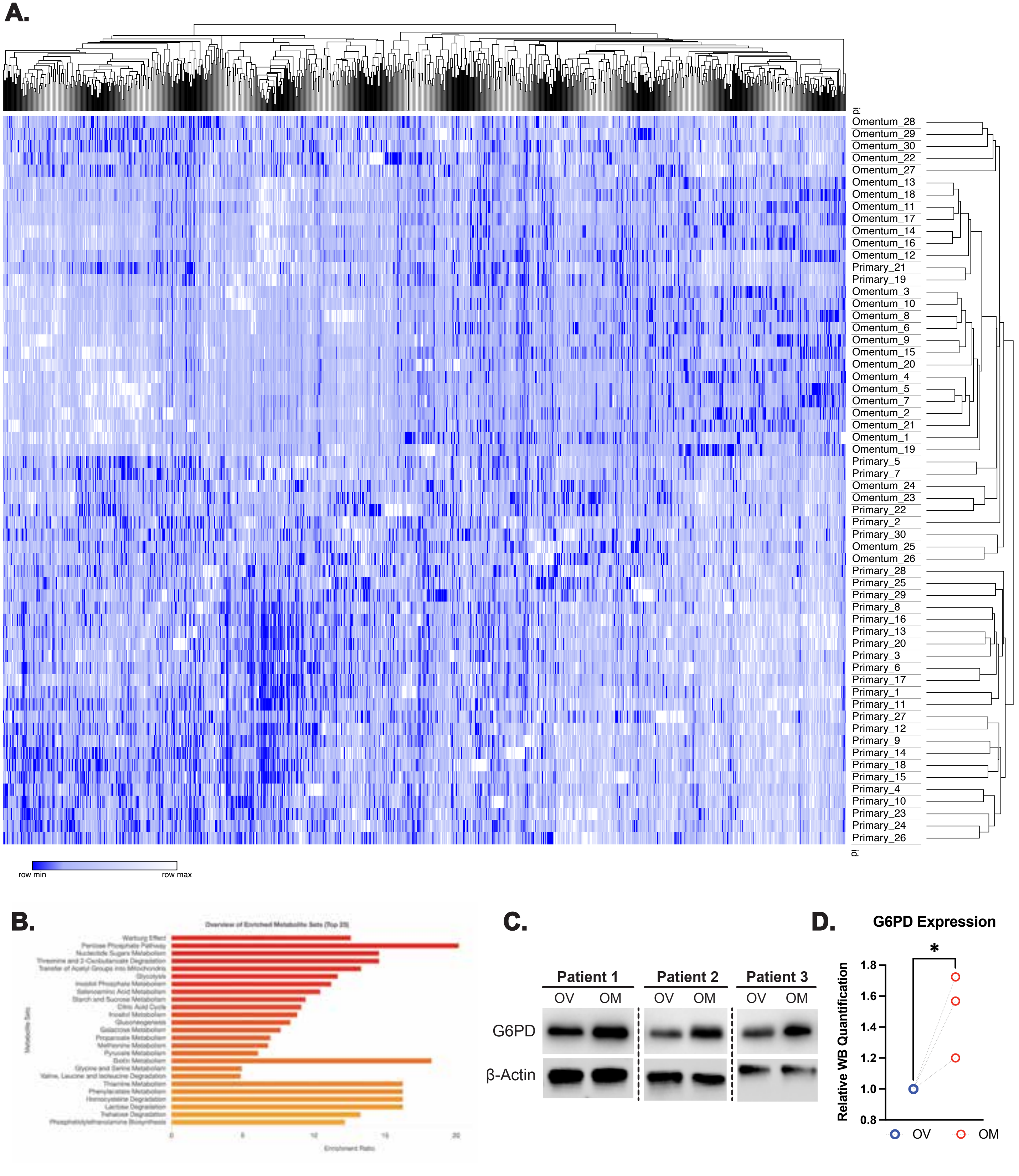

### Supplementary Figure 2

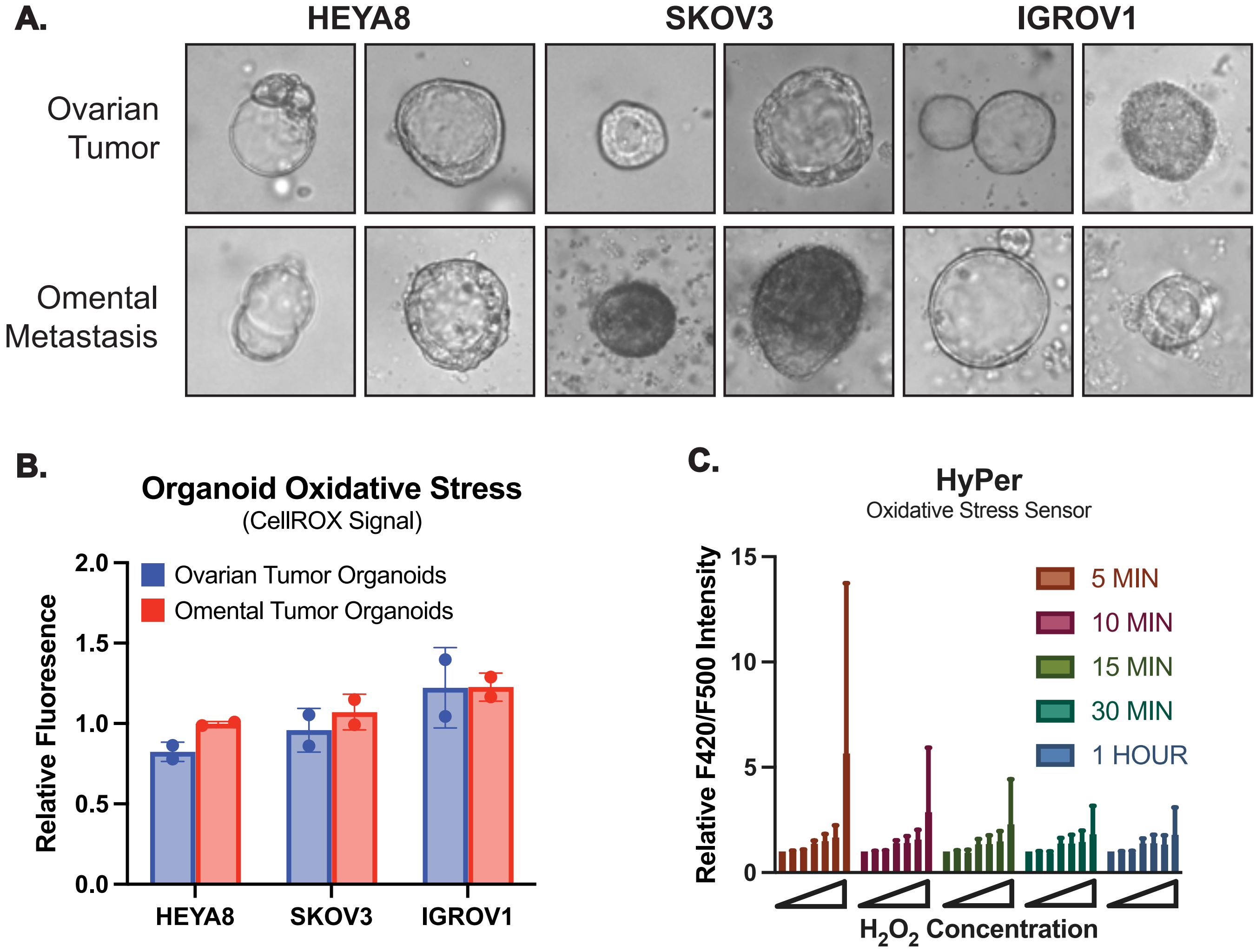

Supplementary Figure 3

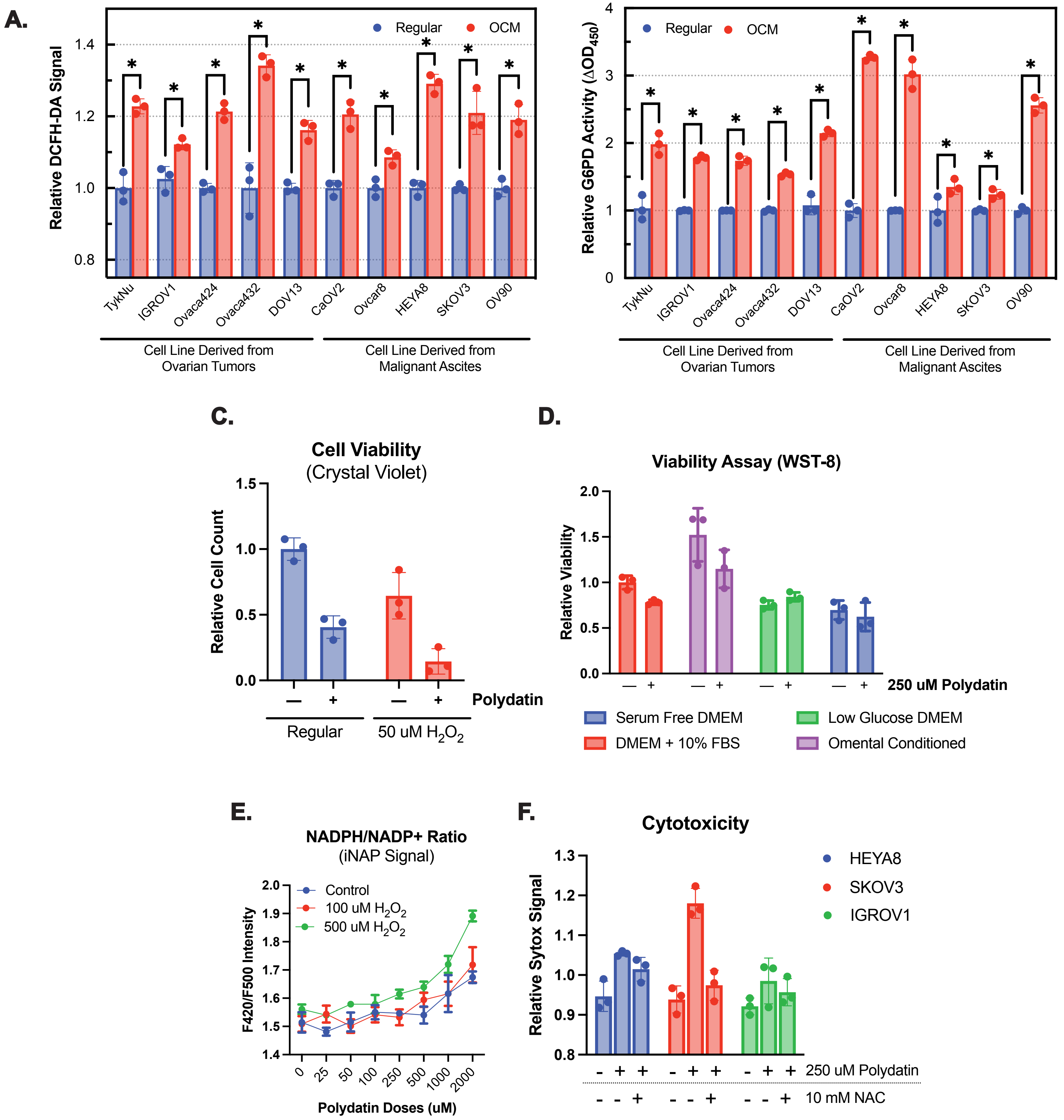

Supplementary Figure 4

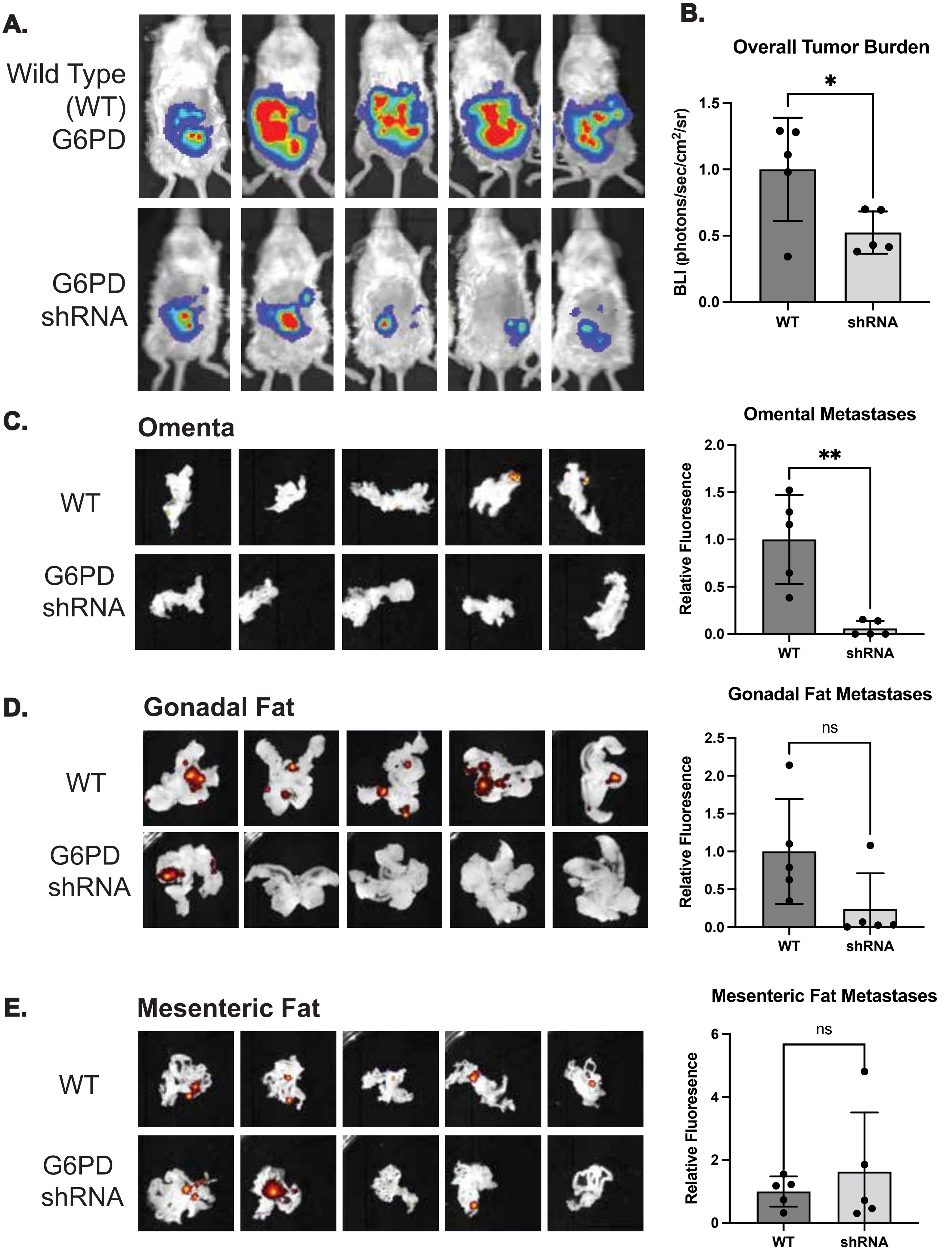

Supplementary Figure 5

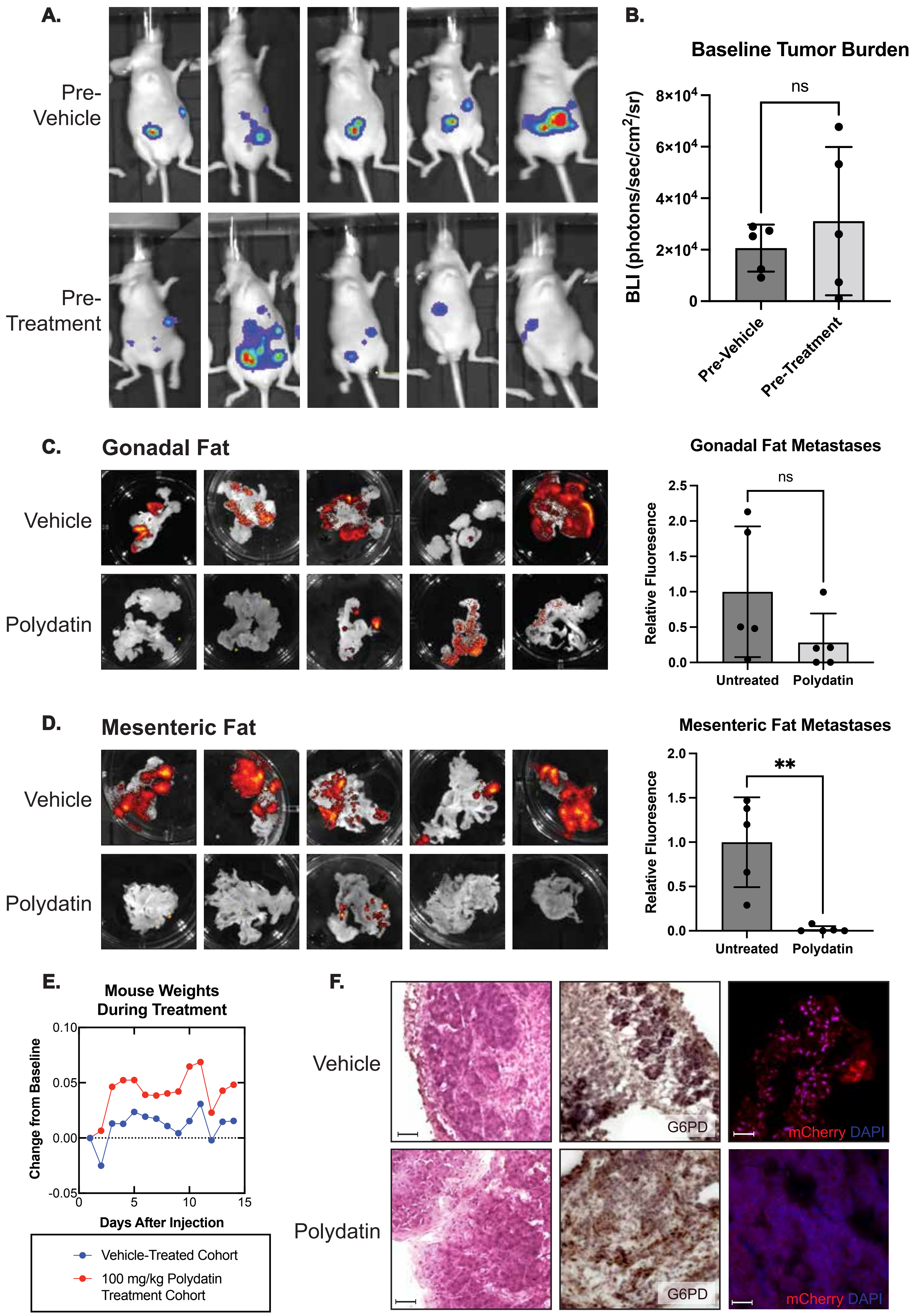

### Supplementary Figure 6

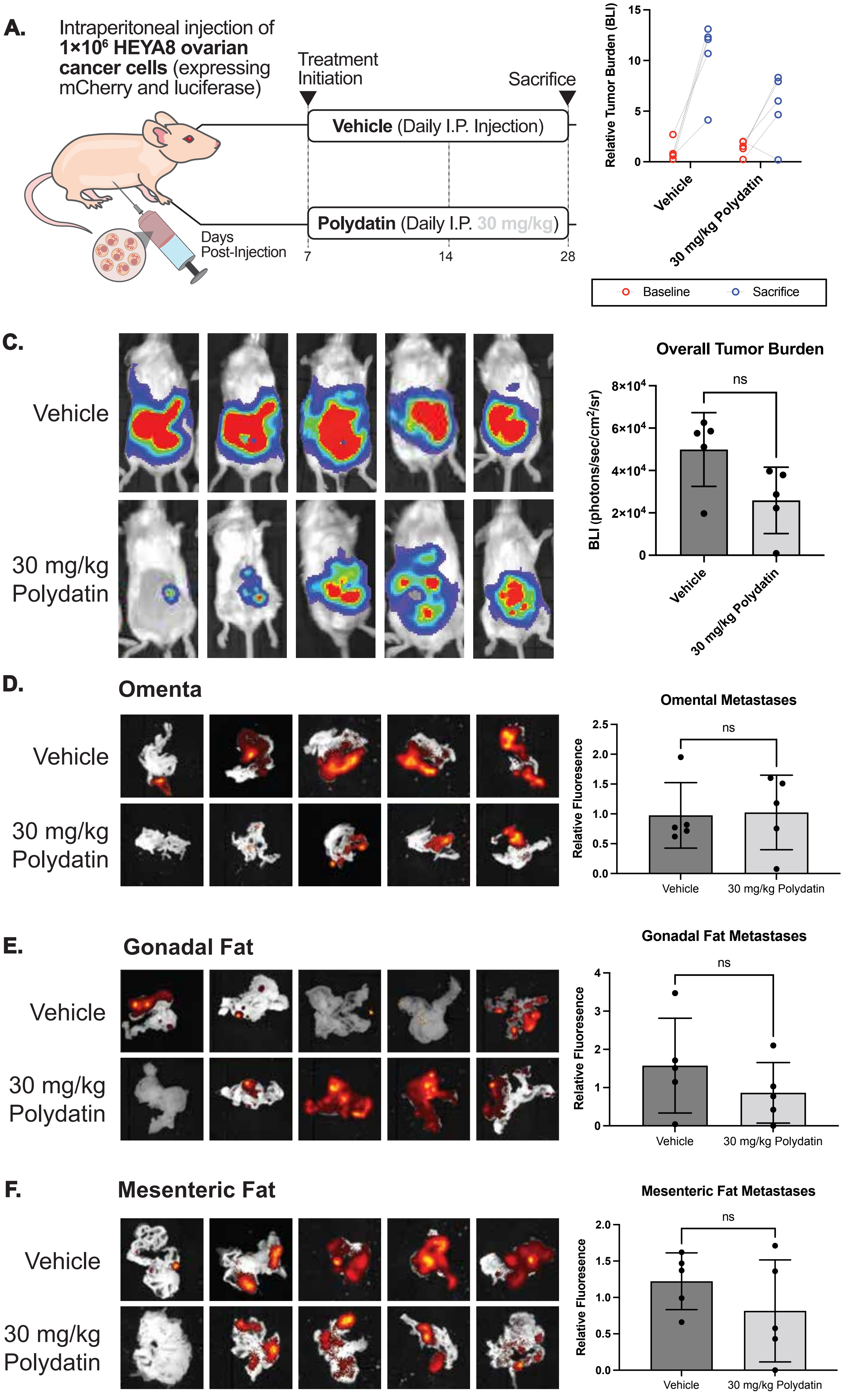
