## Supplemental Table 1 for "G6PD Inhibition Sensitizes Ovarian Cancer Cells to Oxidative Stress in the Metastatic Omental Microenvironment"

| Media Component | Final Concentration |
| --- | --- |
| DMEM/F12 |  |
| GlutaMax | 1x |
| Penicillin/Streptomycin | 100 U/mL |
| 17-B-Estradiol | 10 nM |
| A083-01 | 250 nM |
| B27 (without Vitamin A) | 1x |
| EGF | 50 ng/mL |
| HGF | 10 ng/mL |
| IGF1 | 20 ng/mL |
| N2 Supplement | 1x |
| N-Acetylcysteine | 5 mM |
| Neuregulin I | 10 ng/mL |
| Nicotinamide | 5 mM |
| Noggin | 100 ng/mL |
| R-Spondin 1 | 50 ng/mL |
| SB203580 (p38i) | 1 uM |
| Y-27632 | 10 uM |

**Supplementary Table 1:** OC Organoid Media Composition
